# Not so automatic: Task relevance and perceptual load modulate cross-modal semantic congruence effects on spatial orienting

**DOI:** 10.1101/830679

**Authors:** Daria Kvasova, Salvador Soto-Faraco

**Affiliations:** Center for Brain and Cognition, Universitat Pompeu Fabra, Barcelona; Pompeu Fabra University ICREA, Barcelona

## Abstract

Recent studies show that cross-modal semantic congruence plays a role in spatial attention orienting and visual search. However, the extent to which these cross-modal semantic relationships attract attention automatically is still unclear. At present the outcomes of different studies have been inconsistent. Variations in task-relevance of the cross-modal stimuli (from explicitly needed, to completely irrelevant) and the amount of perceptual load may account for the mixed results of previous experiments. In the present study, we addressed the effects of audio-visual semantic congruence on visuo-spatial attention across variations in task relevance and perceptual load. We used visual search amongst images of common objects paired with characteristic object sounds (e.g., guitar image and chord sound). We found that audio-visual semantic congruence speeded visual search times when the cross-modal objects are task relevant, or when they are irrelevant but presented under low perceptual load. Instead, when perceptual load is high, sounds fail to attract attention towards the congruent visual images. These results lead us to conclude that object-based crossmodal congruence does not attract attention automatically and requires some top-down processing.

## Introduction

Interactions between sensory modalities and their influence on perception and behavior have been convincingly demonstrated over the past decades. For instance, in multisensory contexts, information from different senses influences the deployment of spatial attention (McDonald et al., 2001; Koelewijn et al., 2010; Talsma et al., 2010; Santangelo and Macaluso, 2012). This way, lateralized sounds can produce a shift of attention that facilitates the processing of a visual target presented at that (congruent) location (Spence et al, 1998; McDonald et al., 2000). Even if spatially uninformative, auditory stimuli can enhance the processing of visual events that are temporally congruent (Van der Burg et al., 2008; Van den Brink et al., 2014).

These attention effects by congruent audio-visual stimuli has previously been observed using simple stereotyped objects i.e. Gabor patches, beeps, flashes, by manipulating congruence between low-level attributes such as spatial location or time. Yet, in the real world, multisensory events do not only provide temporally and spatially correlated information but also convey higher-level information about the identity of the object. Like lower level spatio-temporal features, these higher-level attributes can bear congruence relationships, arising from their semantic associations. It is therefore possible that in the natural environment object-based (semantic) relations between sounds and visual events might have an influence on attention orienting. Several recent studies have addressed the role of crossmodal semantic congruence on spatial orienting by investigating how characteristic sounds of objects (musical instruments, vehicles, animals etc) or semantically congruent tactile information can enhance performance in different visual tasks (e.g., Laurienti et al., 2004; Chen and Spence, 2011; Molholm et al., 2004; Pesquita et al., 2013; Iordanescu et al., 2008; Iordanescu et al., 2010; List et al., 2014). However, the results of these studies are mixed, some suggesting that semantic congruence effectively attracts attention and some suggesting that it does not. Because of the great methodological variability between different studies, an important question remains as to what are the conditions in which cross-modal semantic relationships influence orienting behaviors. Answering this question can shed some light on the underlying processes supporting cross-modal semantic interactions.

Nardo et al. (2014) reported that crossmodal semantic congruency between visual events and sounds had no effect on spatial orienting or brain activity (measured with fMRI) when observers watched videos of everyday life scenes. In contrast, another study by Mastroberardino et al. (2015), using static images of objects, reported that attention was oriented toward the image semantically congruent (albeit spatially uninformative) sound presented at the same time. Along similar lines, Iordanescu et al. (2008, 2010) showed that characteristic sounds, even if spatially uninformative, speeded up visual search when consistent with the target object. Conversely to the study of Nardo et al. (2014) which found no effect, Iordanescu et al. and Mastroberardino et al. used simple static images presented in decontextualized search arrays (Iordanescu et al., 2008, 2010). One could argue that this might be the reason for the different result. Indeed, both the dynamic nature of natural scenes and their complexity, have been pointed out as important gaps in the generalization of laboratory research findings to real-world contexts (e.g., Hasson et al., 2010). However, a recent study from our laboratory has addressed these potential explanations by demonstrating that characteristic sounds crossmodally enhance visual search of relevant objects even in complex and dynamic real-life scenes (Kvasova et al., 2019). Therefore, the static stimuli and lack of context in previous studies might not fully account for the difference in the results between previous studies. Here, we investigate whether task relevance might be a factor.

Task-relevance is another possibly important variable that has varied significantly across studies in prior research on cross-modal semantic effects on spatial attention. Iordanescu et al have shown that characteristic sounds, even if spatially uninformative, speed up search times for congruent visual targets (Iordanescu et al., 2008, 2010). In these studies, the visual search array contained four competing stimuli and the visual event was a target itself, the audio-visual object was in this case completely task-relevant. A similar method was applied in Kvasova et al. (2019) expect that the objects were embedded in more realistic video clips, with equivalent results. These studies showed consistent effects of cross-modal semantic congruence when the audio-visually congruent object is relevant for the task at hand (see also Iourdanescu et al., 2008, 2010).

What happens when the audio-visual congruent object is task-irrelevant? In Nardo (2014) participants were asked to freely observe videos without any particular task requirement. In this case, cross-modal semantic relations had null effects on orienting (measured with eye-tracking). In Mastroberardino et al (2015), participants did perform a task, but the audio-visual semantic congruence was putatively task irrelevant. In the cited study, participants were asked to discriminate the orientation of a visual target (a Gabor grating) presented to one side (left or right) of central fixation. However, right before the relevant visual target appeared, a pair of irrelevant images of animals were presented at the corresponding left/right locations where the upcoming targets could appear. What they found is that when a central sound was semantically congruent with one of the two images, then discrimination performance of the visual target presented later at that location improved. Mastroberardino et al. concluded that despite irrelevance to the task, the semantically congruent audio-visual object produced capture, hence summoning attention to that location. Compared to Nardo et al., however, Mastroberardino et al.’s task did not impose a high perceptual load: the presentation the irrelevant animal images was sequential with the relevant visual targets^1^, there was only two of them (always the same two, a cat and a dog), and they were presented at two pre-specified locations. One could even think that the images might have acted as placeholders, and therefore not being completely task irrelevant). In any case, according to the perceptual load theory (Lavie and Tsal, 1994), one would expect processing of irrelevant information under these low perceptual load conditions. Inferring from the results of these very different studies, one might be tempted to conclude that under high perceptual load, cross-modal semantic congruence matters only if it bears some relevance to the task at hand. This is precisely the question that we address in the present study.

Here we aim at investigating how task constraints may modulate the effect of cross-modal semantic congruence on attracting attention. We hypothesized that the effect of audio-visual semantic congruence will emerge when at least one of two conditions apply: the multisensory object (or one of its components) carries some relevance to the current goal or, the multisensory object is irrelevant but presented under low perceptual load. We therefore predict that a semantically congruent sounds speed up the explicit search of a corresponding visual target, but when attention is engaged in another task, and therefore search is not explicit, then this cross-modal semantic effect will wane.

We addressed this question in three experiments using the same set of multisensory stimuli, with the only variations being task relevance and perceptual load. For all the experiments we used audio-visual pairs of common objects (e.g. a picture of a cat and a meowing sound, picture of a phone and a ring tone).

In Experiment 1, we studied the effect of audio-visual semantic congruence on spatial attention when the audio-visual pairs are task relevant. To do so, we aimed at replicating the results of Iordanescu et al. (2008, 2010), where the visual component of the multisensory object was task relevant by explicit instruction. In all cases, participants performed a visual search task for pre-defined visual objects while hearing sounds that could be semantically consistent with the target, consistent with a distracter or not related to any object in the search array. According to these previous results, we expected to find significant effects of cross-modal semantic congruency in the form of shorter search latencies in target-consistent, compared to distractor-consistent trials or neutral trials. Previous findings by Iordanescu et al. (2008, 2010) and Molholm et al. (2004) also show that semantically consistent sounds do not increase the visual salience of a distractor visual objects in the search array. In line with this, no difference in reaction time between distractor-consistent and neutral conditions is expected. This would support (and confirm) that the object-based audio-visual facilitation requires some top-down (goal-directed) processing.

In Experiments 2 and 3, we studied the effect of audio-visual semantic congruence on spatial attention when the audio-visual pairs were not task relevant. In these experiments, an array of visual objects and a sound were presented just like in Experiment 1 (and under the same conditions described). Yet, participants did not have to do any task with this array but were instructed to just wait until this array was replaced with a second visual array composed of “T” letters. Participants searched this second array for an upright “T” amongst inverted “T” s. The variable of interest was whether or not the target in the T search task appeared at a location previously occupied by a congruent audio-visual pair. The difference between Experiments 2 and 3 was perceptual load (low *vs*. high respectively).

According to our hypothesis, the predictions are as follows. If cross-modal congruence triggers automatic orienting, even in task-irrelevant conditions, then we expected to find a search advantage (shorter latency) if targets appeared at the location previous occupied by a congruent multisensory object, compared to when the target appeared away from this location. In the case that these interactions were to occur independently of available processing resources, indicating strong automaticity, then we would expect the effect to survive despite of task irrelevance and high perceptual load (Experiment 3). If orienting toward cross-modal semantic congruence breaks down in Experiments 2 and 3, we will conclude that task relevance is a condition for these interactions. If orienting toward cross-modal semantic congruence breaks down only in Experiment 3, then we will conclude that these interactions may happen even if task irrelevant, as long as perceptual load is low. Either of the two latter outcomes will cast doubts on a strong version of the automaticity hypothesis. Finally, by hypothesis we do not expect the effect of cross-modal semantic congruence to be significant in Experiment 3 but not in Experiment 2. Such a pattern of results should lead to a revision of the initial hypothesis.

## Experiment 1: Replication of Iordanescu et al., 2008, 2010

In Experiment 1 we aimed at replicating the results of the study of Iordanescu et al. (2008, 2010) but with a new set of audio-visual stimuli. We created a visual search task where participants had to look for a target visual object while hearing characteristic sounds that were either consistent with the target of search, consistent with a distractor, or neutral (consistent with neither). We conducted three different versions of the experiment (Experiments 1a, 1b and 1c) with variations in measurement: in Experiment 1a and 1b we measured saccadic search times (Iordanescu et al., 2010) and in Experiment 1c participants gave manual responses (Iordanescu et al., 2008). This was done mainly for the replication purposes and also to ascertain if both types of response are reliable in order to be used in other experiments.

### Experiment 1a: Saccadic responses with aligned audio-visual stimuli

#### Methods

##### Participants

Sixteen volunteers (7 males; mean age 24.56 years, SD = 3.67) took part in the study. They had normal or corrected-to-normal vision, reported normal hearing and were naïve about the purpose of the experiment. All subjects gave written informed consent to participate in the experiment.

##### Stimuli

A set of 20 different images were obtained from free picture data-bases. Images represented tools, animals, transport, etc. (See supplementary materials). All pictures were edited with Adobe Photoshop CC 2015. Each picture was converted to greyscale and scaled to fit within 4.5 ° × 4.5 ° degrees area. All visual stimuli were presented on a gray background. Characteristic audio clips for each of the visual objects were obtained from Freesound.org database (See online supplementary materials). The duration of sounds varied due to differences in their natural durations (M = 660 ms with SD = 130 ms). These differences should not have affected our results since the design of our experiments was counterbalanced (see Procedure). The sounds provided no information about the visual target’s location, were clearly audible and presented via two loudspeakers, one on each side of the monitor, in order to render them perceptually central. On each trial, the sound was either consistent with the target object (*target consistent*), consistent with a distractor object (*distractor consistent*), or not consistent with any of the four objects included in the search display (*neutral*). All of the objects were randomly selected for each trial.

##### Procedure

The experiment was programmed and conducted using Psychopy 1.81 (for Python 2.7) running under Windows 7. An Eyetribe eye tracker (60 Hz sampling rate and 0.5° RMS spatial resolution) with a combined chin and forehead rest was used to control for eye movements.

Participants were sitting in front of a computer monitor 22.5’’ (Sony GDM-FW900) at a distance of 77cm. In order to start each block of the experiment, participants pressed the space bar. Each trial started with the fixation cross that lasted for 2000 ms. Then a cue word was printed on the screen indicating the target of the visual search for that trial (Figure 1). After 2000 ms, a cue word disappeared, and a search display appeared. Every trial of the experiment contained a display with 4 black and white pictures of visual objects that were placed in the four quadrants at 4.7° eccentricity. One of these four objects was a visual search target and the rest three were distractors. Visual display with objects appeared simultaneously with the sound that followed one of the three experimental conditions (*target-consistent, distractor-consistent* or *neutral*).

**Figure 1.**
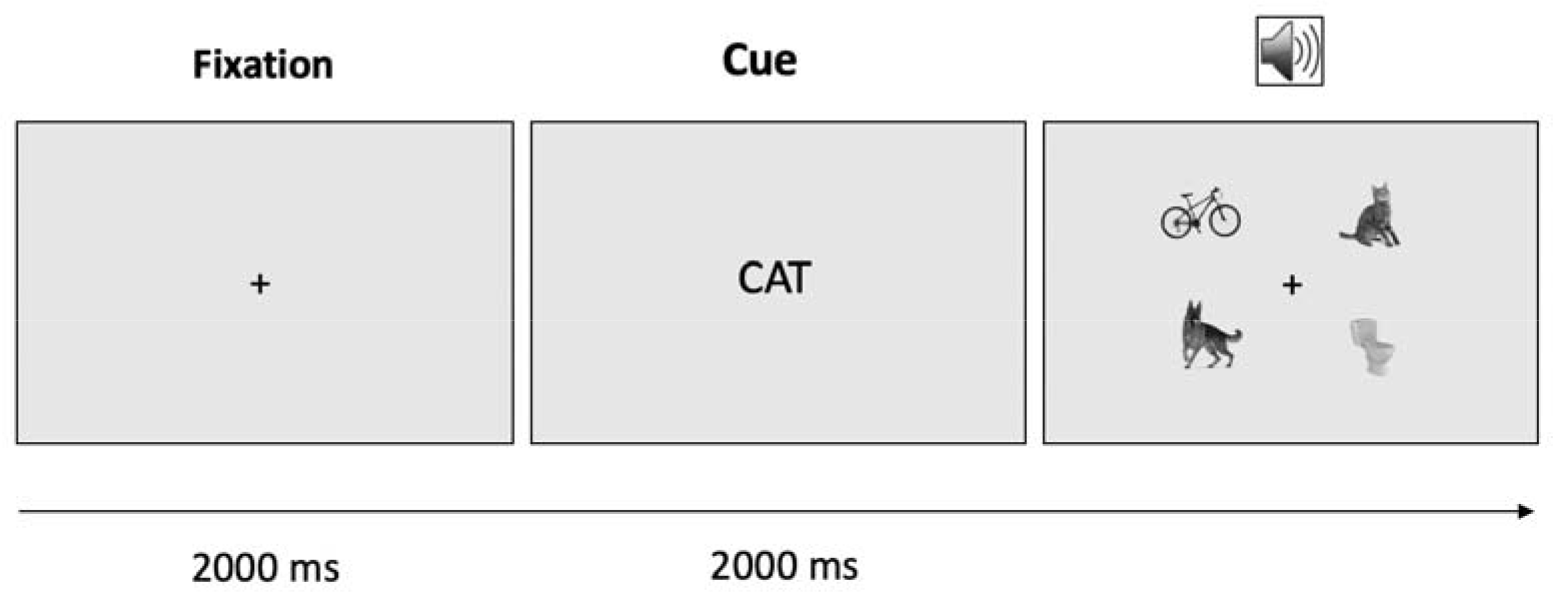
The sequence of event was identical for the Experiments 1a, b and c. First participants were asked to stay fixated in the central cross. The fixation cross was followed by the presentation of a cue word. Then search array of 4 objects appeared together with the sound that in the Experiment 1a was synchronous to the visual onset and in the Experiments 1 b, c preceded the pictures for 100 ms. In the Experiments 1 a, b participants had to look at the target object as fast as possible. Once the gaze was detected inside of the quadrant with target the trial was abrupted and the new trial began. In the Experiment 1c participant had to maintain central fixation throughout the whole trial and press as fast as possible one of the four keys that corresponded to the location of visual target. In contrary to the Experiment 1 a and b, in the Experiment 1c visual objects were presented only for 670 ms. Participant could respond during these 670 ms, otherwise the question mark appeared and stayed until participant presses the response key.

Participants were instructed to look as fast as possible at the visual target. Visual search performance for each subject and condition was determined by the mean Saccadic search time (SST). Once eye gaze entered the quadrant with visual target the trial automatically finished, and the new trial begins. SST was calculated from the beginning of the appearance of the visual search display until the moment the left eye gaze position reached the region of the target. Target object was presented in every trial. The experiment consisted of 4 blocks in total, each block contained: 20 target-consistent, 20 distractor-consistent and 20 neutral trials.

#### Results

We ran a repeated measures ANOVA on mean search times, with subject as the random effect and condition as the factor of interest. The analysis showed that main effect of condition was not significant (F(2,30)=0.4; p=0.67). Saccadic search time was not significantly faster in the target-consistent-sound condition (M = 501 ms) compared with both the distractor-consistent-condition (M = 500 ms), t(15) =0.29, p = 0.388 and neutral condition (M = 497 ms), t(15) = 1.05, p = 0.154. Neither the difference was found between the distractor-consistent and neutral conditions, t(15) =0.56, p = 0.29 (Figure 2). Thus, in the Experiment 1a characteristic sounds did not speed up gaze towards the visual target.

**Figure 2.**
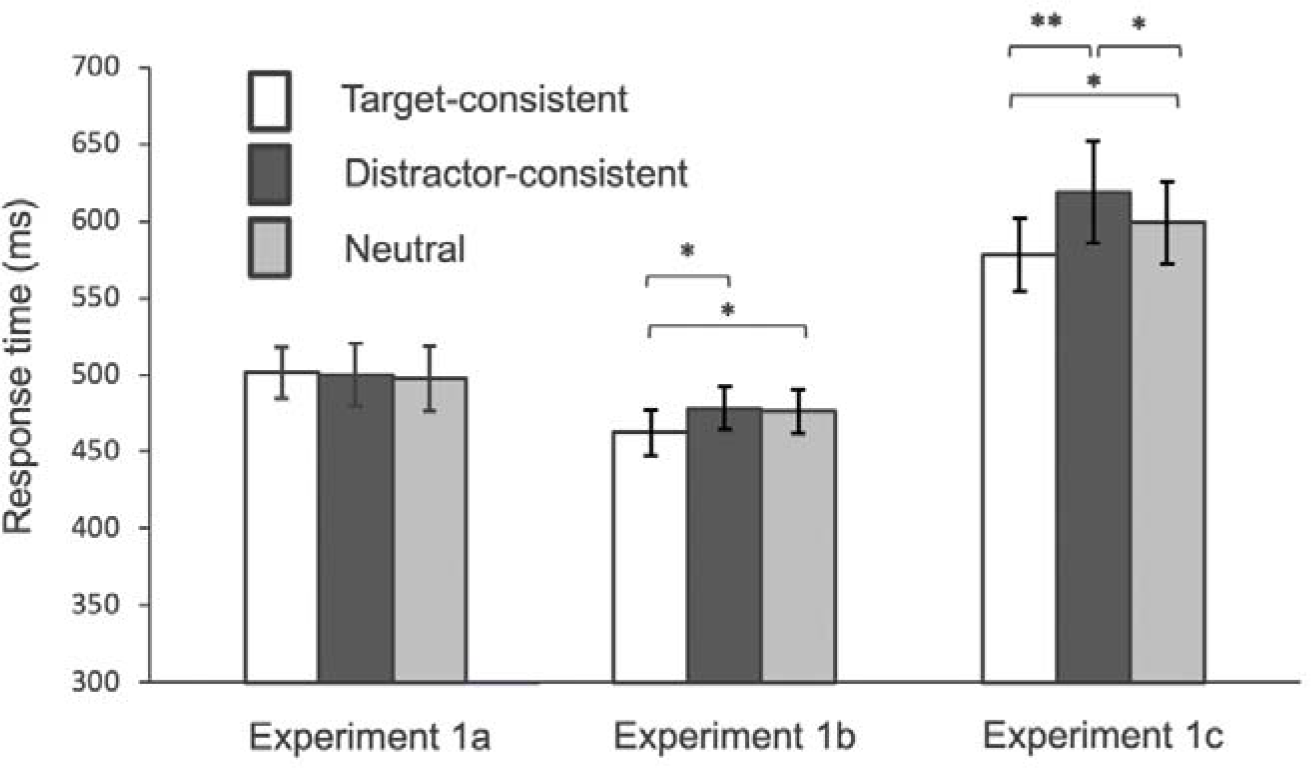
Visual search average reaction times towards a target and error rates were plotted in the *target-consistent* sounds, *distracter-consistent* sounds and *neutral* sounds conditions in Experiment 1a (Saccadic search times, SOA0ms), Experiment 1b (Saccadic search times, SOA100ms) and Experiment 1c (Manual response times, SOA100ms). Error bars indicate the standard error. Asterisks indicate significant difference between conditions (1 asterisk for p-value less than 0.05, 2 asterisks for p-value less than 0.01)

The results of this experiment fail to replicate the cross-modal semantic effect, a finding established by several previous studies. One of the reasons for the null result might lie in the nature of processing of complex sounds. Meaningful sounds take a certain (and variable) amount of time to identify given that information needs to be integrated over some hundreds of milliseconds (e.g. Cummings et al., 2006). On the other hand, less time is necessary to access the meaning of visual information in comparison to (Kim et al., 2014; Weatherford et al., 2015). In particular, semantic information can be accessed from visual stimuli within the first 100ms (for a review, see Potter, 2014), whereas processing of the meaning of a complex naturalistic sound can require more time due to the temporal nature of the information (according to some review, approximately 150 ms after onset, Murray and Spierer, 2009). For this reason, the temporal window of audio-visual integration for complex sounds is not the same as for simple artificial sounds (Vatakis and Spence, 2010). Here, because saccades are fast, it might be the case that there was no sufficient integration time for the meaning of the sound to influence visual processing before response. Following the same logic as previous laboratory studies that used complex sounds and visual events, we decided to advance the presentation of sounds by 100ms in Experiment 1b (Vatakis and Spence, 2010, for a review; Knoeferle K. M., Knoeferle P., Velasco and Spence, 2016, Kvasova et al, 2019, for a similar procedure). In Experiment 1c, we used the same procedure but changed the type of response: instead of saccadic search we used manual response.

### Experiment 1b: Saccadic responses with offset sounds

All apparatuses and stimuli in Experiment 1b (see *Experiment 1a Methods*) were identical to those used in Experiment 1a, except that the sound preceded the onset of visual search array for 100 ms (stimulus onset asynchrony SOA 100ms). For this purpose, we recruited an additional group of 16 participants (7 males; mean age 24.56 years, SD = 3.67).

#### Results

The analysis returned a significant main effect of condition (F(2,30)=4.14; p=0.025). Further, we tested the differences between conditions using one tail t-test with Holm-Bonferroni correction for multiple comparisons (Ludbrook, 1998). The analysis showed that saccadic search time was significantly faster in the target-consistent sound condition (M = 462 ms) compared with both the distractor-consistent (M = 478 ms), t(15) =2.51, p = 0.012, Cohen’s d=0.27 and neutral condition (M = 476 ms), t(15) = 2.55, p = 0.011, Cohen’s d=0.23, see Figure 2). All these results survived the correction for multiple comparisons. Thus, in Experiment 1b we have shown that during visual search task characteristic sounds, when presented 100 ms in advance, speeded gaze responses towards semantically congruent visual targets. This successfully replicates previous results and establishes the cross-modal semantic effect on visual search. In addition, we compared search times in distractor-consistent and neutral conditions. The additional prediction stated that if audio-visual semantic consistency has an impact only in a goal-directed way then distractor-consistent sounds should not slow down performance compared to other not related sounds. Post-hoc t-tests showed the lack of difference in saccadic search time between the distractor-consistent and neutral conditions, t(15) =-0.902, p=0.381.

### Experiment 1c

An additional group of 16 participants was recruited for this experiment (7 males; mean age 24.56 years, SD = 3.67). The experiment was programmed and conducted using the MATLAB 8.2-R2013b running under Windows 7. The procedure in the experiment 1c was adapted for the use manual response instead of gaze responses. The details of the procedure are as in Experiment 1b, except for the following differences: In experiment 1c participants had to maintain visual fixation throughout the whole trial, and were instructed to press one key out of four possible (1, 7, 9 and 3 keys of the number pad of the keyboard) corresponding to the location of the target (instead of gazing at the target). Visual search performance for each subject and condition was determined by the mean reaction time (RT) of manual responses. RT was calculated from the beginning of the visual search display until the moment the moment subject pressed the response key.

#### Results

We ran a repeated measures ANOVA on mean RTs (over correct responses), with subject as the random effect and condition as the factor of interest. The analysis showed a significant main effect of condition (F(2,30)=7.7; p=0.002). We found that reaction time was significantly faster in the target-consistent-sound condition (M = 578 ms) compared with both the distractor-consistent-condition (M = 619 ms), t(15) =3.05, p = 0.004, Cohen’s d=0.36 and neutral condition (M = 599 ms), t(15) =2.51, p = 0.012, Cohen’s d=0.19 (Figure 2). All comparisons survived the Holm-Bonferroni multiple comparison correction. Furthermore, reaction time was slower in distractor-consistent than in neutral conditions, t(15) =2.36, p = 0.016, Cohen’s d=0.17. This last result suggests that distractor objects may also attract attention when congruent with the sound, even if these audio-visual events are not relevant to the current goal. This result was not expected, by comparison to previous results in the literature (and with the results of saccadic responses in Experiment 1b). We will come back to this in the General Discussion section. All in all, the results of experiments 1b and 1c allowed us to show that semantically consistent sound attract attention to the visual object when audio-visual event is relevant to the task, i.e. visual search. This replicates the finding from Iordanescu. (2008, 2010) and also Knoeferle et al. (2016) and Kvasova et al. (2019). Because the effect of crossmodal semantic congruence was found only when sound was presented 100ms before the visual stimuli we decided to use SOA100ms in all the following experiments. Also, the effect size was larger when using manual versus eye responses. Therefore, we used manual responses for the subsequent experiments.

## Experiment 2

In the previous experiments we found that when audio-visual pair is relevant to the current goal of the task, semantic congruence has an effect on search. As the next step of our study addressed whether audio-visual semantic congruence attracts attention even when the audio-visual pair is not relevant to the task. Here participants saw the same arrays as in experiments 1A-C, containing images and characteristic sounds of common objects (same conditions), only this time these arrays were completely task-irrelevant. Instead, subjects were asked to wait until the array transitioned into a new display composed of a set of letters T, and then perform a search task for an upright T amongst rotated Ts. In some of the trials the target T appeared at the same spot where the visual object congruent with the sound had been presented before. If audio-visual semantic congruence attracts attention to its location, then we would expect benefits in visual search if the target of the new away falls at that location. Because audio-visual events are task irrelevant, this would mean that crossmodal sematic congruency is able to attract attention in an automatic manner.

### Methods

#### Participants

Fifteen volunteers (6 males; mean age 26.32 years, SD = 4.15) took part in the study. They had normal or corrected-to-normal vision, reported normal hearing and were naïve about the purpose of the experiment. All subjects gave written informed consent to participate in the experiment.

#### Stimuli and Procedure

For this experiment we used the same set of 20 pictures of common objects and their corresponding sounds as in Experiment 1. We presented sounds 100 ms before visual onset similarly to Experiments 1 b & c. In the beginning of the trial participants were presented with the cross in the middle of the screen for 500 ms and were instructed to maintain visual fixation on it (Figure 3A). Then 4 pictures of common object together with a sound were presented for 670 ms. The sound was either consistent with the object that was located in the quadrant where the following target of search will appear (*cued*); consistent with the object located in one of the other 3 quadrants (*uncued*), or not consistent with any of the four objects (*neutral*). After that, pictures faded out gradually and the search array started to appear on top of it. This transition lasted for 150 ms until pictures of objects completely disappeared and the search array was clearly visible. The transition was used in order to avoid abrupt changes that might induce the reorientation of attention (Remington et al., 1992). The search array contained 16 ‘T’ letters. Participants searched for an upright “T” within inverted “Ts” and were instructed to press a response key as fast as they could when they found the target or withhold response if there was no target (filler trials).

**Figure 3.**
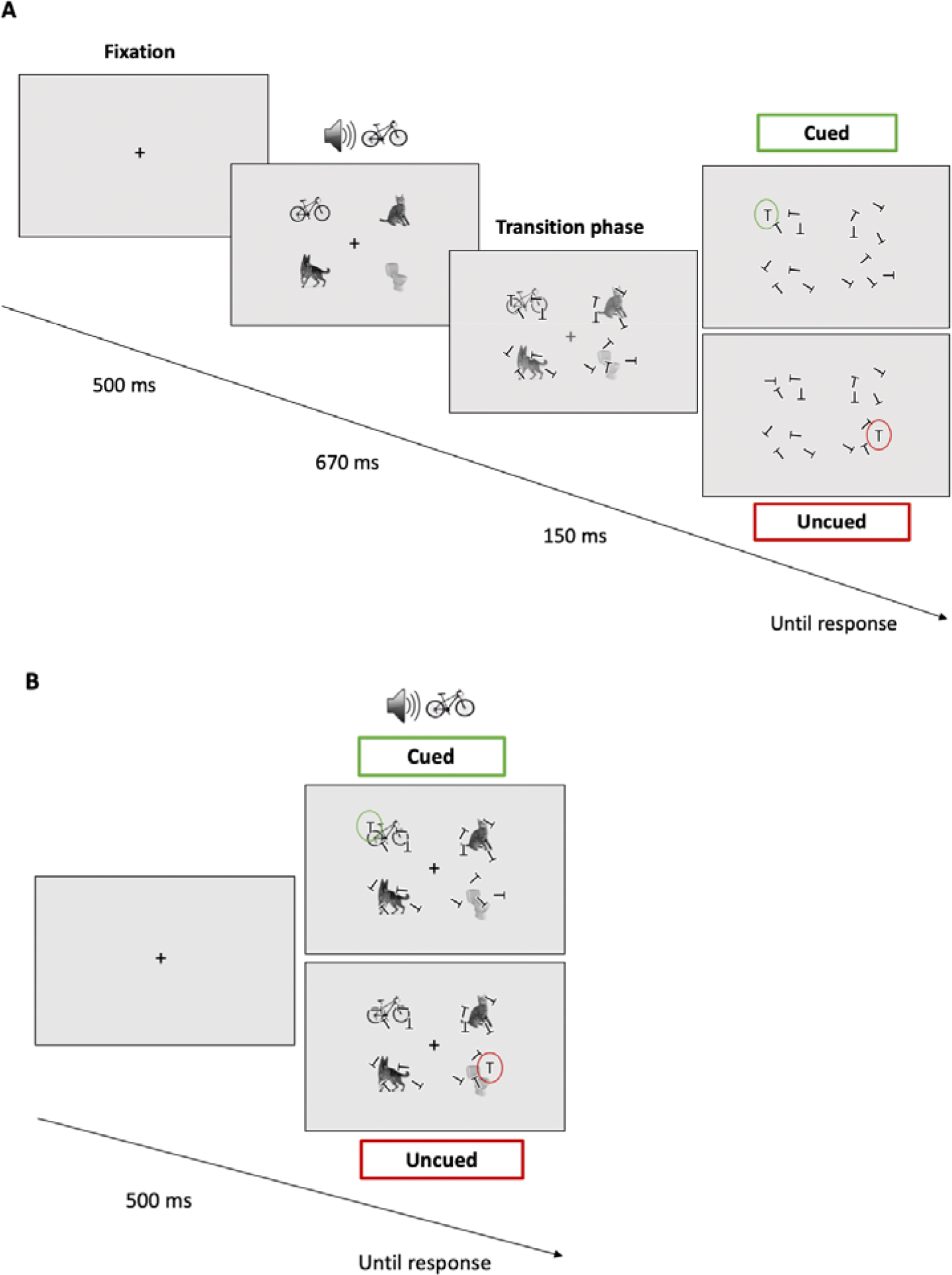
A) Sequence of events in the Experiment 2. At the beginning of the trial fixation cross appeared for 500 ms. 4 pictures of objects were then presented with the centrally presented sound for 670 ms. Sound advanced the presentation of the pictures for 100 ms. Then the trial continued into the transition phase for 150 ms during which the pictures gradually disappeared and search screen with inverted ‘T’ letters appeared. Search screen stayed until participant responds. B) Experiment 3 included two types of trials: with low (A) and high perceptual load (B). In the high perceptual load trials participants viewed pictures with sounds together with search array of inverted ‘T’ letters. Search screen stayed until participant responds.

The experiment consisted of 4 blocks of 200 trials each. In total 800 trials: 160 trials with no target (filler trials), 120 cued, 160 neutral and 360 uncued trials. By the low presence of cued trials (15% of the total) we disincentivized the strategy of anticipating targets where the previous audio-visual congruent event was located, which could artificially generate the result we expected.

This procedure was adapted from the study of Mastroberardino et al. (2015). However, we did several important modifications. In the study of Mastroberardino et al. (2015) authors primed location of the upcoming relevant visual events with 2 object images (always a cat and a dog) that repeated throughout the experiment and were always at the same two locations. Because of this, and the strictly sequential presentation of the images and the visual targets, the perceptual load and the competition for processing resources between stimuli was relatively low. Also, since the initially irrelevant visual objects consistently market the two positions where visual targets could appear could have become relevant to the task. In Experiments 2 and 3 of the current study, displays contained 4 pictures of common objects selected from a set of 20 different objects randomly chosen in every trial. In Experiment 2 we used sequential presentation of events (just like in Mastroberardino et al., 2015), whereas in Experiment 3 (described below) the image array and the search array appeared concurrently. Finally, in the present experiments the target task required visual search with 16 T letters that were equally distributed within the area where previous 4 pictures appeared. This helped us to avoid the role of pictures acting as placeholders and therefore minimize possible relevance to the task which was important to our research question.

### Results

We anticipated that, if crossmodal semantic congruence attracts attention despite being irrelevant to the task then search time in the cued condition should be faster than in uncued or neutral. ANOVA returned a significant main effect of condition, F(2,28)=5.56 p=0.009. The analysis showed that average RTs in the cued condition (M = 893 ms) were significantly faster than in the uncued (M = 955 ms), t(14)=3.96, p=0.0003, Cohen’s d=0.34 or neutral condition (M=937 ms), t(14) =2.02, p=0.031, Cohen’s d=0.24 (Figure 4A). All these comparisons are one tail (given the directional hypothesis) and survived the multiple comparison correction using Holm-Bonferroni.

**Figure 4.**
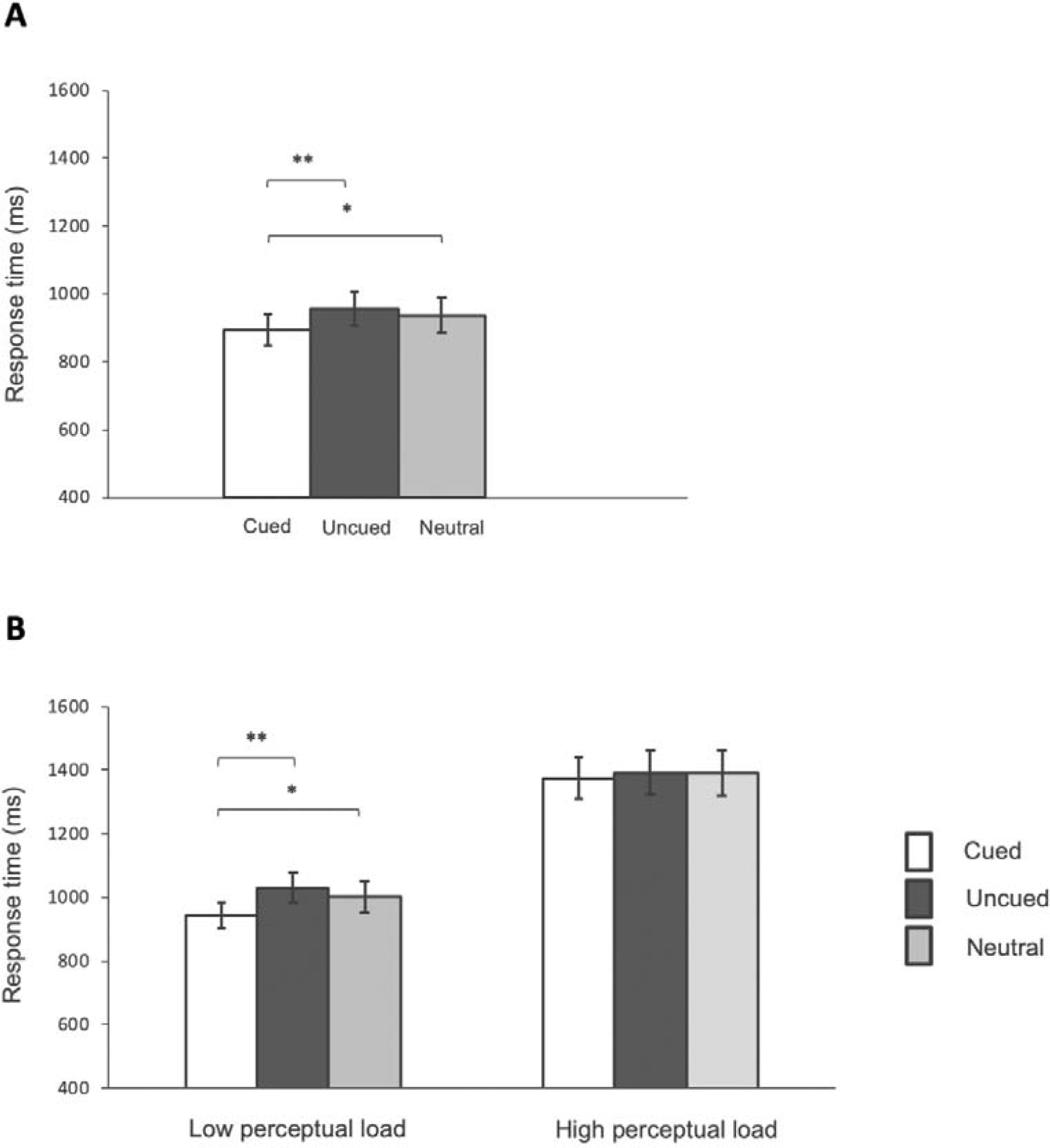
**A) Experiment 2:** Visual search reaction times towards a target and error rates were plotted in the *cued* (white), *uncued* (black) and *neutral* (grey) conditions. **B) Experiment 3:** Visual search reaction times towards a target and error rates were plotted in the *cued* (white), *uncued* (black) and *neutral* (grey) conditions and separated in two plots. Left plot represents performance in the same three conditions as in the Experiment 2 in the low perceptual load and right plot in the high perceptual load trials. In both experiments error bars indicate the standard error and asterisks indicate significant difference between conditions (1 asterisk for p-value less than 0.05, 2 asterisks for p-value less than 0.01)

Our results demonstrate that semantically consistent sounds guide attention to its corresponding visual object despite the fact that the audio-visual events are irrelevant to the current task. Although we introduced several important changes in the design to increase uncertainty and irrelevance of audio-visual events to the task, we still have observed the effect of crossmodal semantic congruence on orienting, similarly to the study of Mastroberardino et al. (2015). However, despite we assumed task-irrelevant design, the presentation of events in the Experiment 2 was still sequential, since sounds and visual objects were always presented before the actual task. Therefore, the perceptual load in this study was relatively low, liberating resources required for the perceptual processing of pictures and sounds that despite being irrelevant to the current goal appear to the participant in the moment when no other events took place. If cross-modal semantic interactions are strongly automatic, then we would expect the effect to survive not only task irrelevance, but also high perceptual load. This was tested in Experiment 3.

## Experiment 3

In Experiment 3 we preserved the irrelevance of audio-visual events to the task, but we introduced the additional differentiation between high and low perceptual load. The task was the same as in the Experiment 2 (see *Experiment 2. Methods*) and the presentation of sounds followed the same conditions. However, the perceptual load in this experiment was high since all objects and sounds were presented together with the array of “T” letters upon objects (Figure 3B). To be able to directly compare the effect of perceptual load we included both types of trials *high* vs *low* (intermixed within blocks). An additional group of twenty participants was recruited for Experiment 3 (7 males; mean age 24.45 years, SD = 3.05). The experiment was divided in 8 blocks of 200 trials: 2 types of load (low and high) × (20 no target) + (15 cued) + (20 neutral) + (45 uncued). Hence, this experiment contained a total of 1600 trials (160 no target, 120 cued, 160 neutral, and 360 uncued, per each load condition).

### Results

We run a repeated measurements ANOVA separately for high and low perceptual load conditions. The analysis of the low perceptual load data returned a significant main effect of sound condition (F(2,38)=7.56; p=0.002). Further comparisons showed that in the trials with low perceptual load average RTs in the cued condition (M = 948 ms) were significantly faster than in the uncued (M = 1028 ms), t(14)=3.57, p=0.001, Cohen’s d=0.40 or the neutral condition (M=1005 ms), t(19)=2.41, p=0.013, Cohen’s d=0.30 (Figure 4B). This replicates the effects found in Experiment 2. All significant effects survived Holm-Bonferroni correction for multiple comparisons. Instead, in the high perceptual load no effects of sound were found F(2,38)=0.36, p=0.7. Reaction time was not significantly faster in cued (M = 1375 ms) and uncued trials (M = 1393 ms), t(19)=0.72, p=0.24. Neither the difference between cued and neutral (M = 1391 ms) conditions resulted significant t(19)=0.58, p=0.28. These results demonstrate that audiovisual congruent events can summon attention even when task irrelevant, but only in low perceptual load condition. Attention capture by cross-modal semantic congruence does not survive high perceptual load.

## Discussion

We addressed whether, and under which conditions, semantic congruence between sounds and visual objects attracts visual spatial attention. We manipulated task relevance of the audio-visual object and perceptual load. The findings to emerge from the experiments presented in this study show that audio-visual semantic congruence can help improve performance when searching for objects that are relevant to the current task goal. When task-irrelevant, the extent to which audio-visual congruence may attract visual attention is limited by perceptual load.

In Experiment 1 (Experiments 1B and 1C) characteristic sounds speeded up search times for the semantically corresponding visual target in a visual search task. This result is in agreement with the idea that cross-modal semantic congruence can attract spatial attention and confirms prior results (Iordanescu et al, 2008, 2010; Knoeferle et al., 2016; Kvasova et al., 2019). In Experiment 1B, distractor consistent sounds did not slow down responses compared to neural sounds, suggesting that audio-visual congruence benefits goal-directed processes, but not the processing of other potential objects. However, in Experiment 1C we found that distractor-consistent sound slowed down search latencies in comparison to neutral sounds as well. This result is against the hypothesis and rather suggests that despite the irrelevance to the current goal semantically congruent audio-visual distractor attracted attention.

In Experiment 2, we measured search times in a visual array unrelated to the audio-visual objects, presented right after. The results showed that, when perceptual load is low, search times benefit if a visual target appears at a location previously occupied by an audio-visually congruent but task-irrelevant object. This finding suggests that cross-modal semantic congruence can attract spatial attention even if not bearing any particular relevance to the person’s task. Therefore, the previous notion that semantic audio-visual enhancements occur only in a goal-directed task-relevant manner is not fully supported (Molholm et al., 2004; von Kriegstein et al., 2005; Iordanescu et al., 2008, 2010). The results of Experiment 2 are in line with the study of Mastroberardino et al. (2015) showing that, despite being irrelevant to the task, crossmodal semantic congruence can still attract attention.

In Experiment 3, we used the same task as in Experiment 2, but used two different perceptual load conditions. In the low load condition, an exact replication of Experiment 2, we found the same results: Task irrelevant audio-visual congruence attracted attention. In the high perceptual load condition, we found that that effect of task-irrelevant semantic congruence just vanished.

What consequences do these results have to interpret prior findings? We believe that the effect of perceptual load might help explain why some studies find task-irrelevant effects of semantic congruence, and why some others do not. For example, this could help explain the unstable effect of distractors in our Experiments 1B and 1C. Contrary to ours, Mastroberardino found crossmodal semantic effects on spatial orienting only for difficult visual targets, in one of the two experiments they reported. The authors suggested that probably, contrary to Iordanescu et al. (2008, 2010) both valid and invalid audio-visual events acted as distractors to the current task. However, we believe that difficulties in finding a stable effect could be rather explained by abrupt change between presentation of audio-visual events and task display. This sudden switch between displays might induce the reorientation of attention (Remington et al., 1992) that further vanished all the cueing effects of semantically congruent audio-visual pair. We believe that the transition phase between displays used in the current study helped to maintain the attention on the location where previous congruent event appeared.

In sum, the results of Experiment 2 and 3 have shown cross-modal semantic congruence can attracts spatial attention even if not bearing any particular relevance to the current task. This pattern of results would suggest that audio-visual congruent objects do have a tendency to attract attention in an automatic manner. However, we also found that perceptual load might act as a limiting condition to this automatic tendency for congruence effects. This speaks against a strong automaticity account of cross-modal semantic interactions.

In order to conclude that audio-visual semantic interactions are fully automatic it would be necessary to demonstrate that the effect appears in task-irrelevant conditions and survives when attention is compromised by high perceptual load. The results of Experiment 3 suggest otherwise. Under high perceptual load when the number of items for processing is high and therefore the amount of resources is exceeded, the effect of audio-visual semantic consistence disappears when task irrelevant. This means that audio-visual semantic congruence necessitates from some top-down regulation in order to guide attention, above and beyond any fast, bottom up cross-modal integration process. This cross-modal interaction can be triggered even in the absence of a particular goal, as long as sufficient processing resources are left available. However, if so, the crossmodal sematic effect should be observed in the high perceptual load condition in Experiment 3. Instead, we found that when attention is fully engaged in different task semantically congruent audio-visual event does not attract attention. Therefore, we believe that semantic-based audio-visual integration requires some attention.

Even if one cannot conclude on automatic cross-modal effects, one interesting question is still open about what is the mechanism of audio-visual semantic interactions. One might think that this pattern of results could be based on the well-known effect of semantic priming, without the need to invoke a different process of fast semantic integration across modalities. Cross-modal facilitation by semantic priming could be explained by the fact that semantic associations across modalities established via prior experience tend to be reinforced. When information in one sensory modality recalls semantic representations, it creates expectation in other modalities which enhances recognition of the upcoming information that is congruent (e.g., Parise and Spence, 2009). Previous studies have demonstrated priming effects across modalities, however, the effect was observed using asynchronous presentation of auditory and visual events and null effect for synchronous or nearly synchronous presentation (Chen and Spence, 2011; 2013). Previous studies generally suggest that cross-modal semantic priming appears when cues (e.g., sounds) are presented prior to targets (e.g., the visual stimulus) (Dehaene et al., 1998; Costello et al., 2009). In our current study, however, consistent sounds were presented only 100 ms before the visual onset, which lead us to suggest that this effect might be caused by a different mechanism than traditional crossmodal semantic priming. Such mechanism would have to be based on interactions between quickly accessed auditory and visually identity information. Possibly, crossmodal semantic effects happen as well due to automatic audio-visual processing based on semantic information. This notion is supported by the findings in the recent study of Cox et al. (2015). Authors showed that synchronously presented semantically congruent sound boosted visual below threshold image into the awareness during continuous flash suppression (CFS). No effect was found when sounds preceded the image. The authors suggested that these cross-modal semantic effects are due to automatic audio-visual processes, rather than traditional semantic priming.

Despite the interaction mechanisms alluded to in the discussion above provide potential accounts for our effects, the actual impact of traditional priming mechanisms is still difficult to assess. For example, in the previous studies where presentation of visual stimuli was very brief (e.g. 27 ms in the study of Chen & Spence, 2011), synchrony manipulations may have been effective to attribute the effect of priming, which are supposed to unfold in time. In our case, the duration of stimulus presentation was relatively long (approximately 660 ms for sounds and 670 ms for pictures). Given that the temporal overlap between auditory and visual stimuli was large, the effects of semantic priming may still occur over the time-course of synchronized events. However, the methods of the current study do not allow us to conclude on the mechanisms of audio-visual semantic interactions. More studies should be conducted in order to address this question.

Given the results observed in this (and prior) experiments, one important question are the implications for real-life scenarios. Object-based enhancements occur consistently for task-relevant objects and might occur even when task irrelevant but only under favourable, low load, conditions. However, like in previous demonstrations of the same principle, experiments have typically used rather artificial settings: stimuli are presented under relatively low perceptual load, and without any meaningful context and ecological validity (Iordanescu et al., 2008; 2010; Mastroberardino et al., 2015). This is unlike real world conditions, where functional relationships and statistical regularities between objects (forks are often seen next to dishes), or between an object and its context (cars are rarely part of a submarine scene), are of great importance. Previous visual-only studies have already made a point about the differences in how attention is distributed in naturalistic, real life scenes compared to simple artificial search displays typically used in psychophysical studies (e.g., Peelen and Kastner, 2014, for a review; Henderson and Hayes, 2017).

Recently, the importance of studying multisensory interactions in realistic environments has been highlighted (e.g. Soto-Faraco et al., 2019; Matusz et al., 2019). One particularly relevant point refers to the interaction between these multisensory processes and attention, given that in realistic contexts, perceptual load tends to be high, compared to the idealised conditions of the current (and previous experiments). According to our results, high perceptual load leads to a decrement in the effectivity of crossmodal congruent events to attract attention (see also, Lunn et al. 2019). This could mean that the incidence of these effects in real life contexts could be limited.

As discussed above, another important aspect of real-world scenes, compared to the artificial displays used here, is contextual information. Whereas in artificial search displays the different elements and their location do not provide any particular constrain on each other, naturalistic scenes are precisely defined by learned relationships that have an impact on object identification (see Peelen and Kastner, 2011). Under this light, in real-world settings that contain information from multiple sensory modalities, semantic relationships might be especially important for orienting. Further studies to understand the limits of crossmodal semantic effects and how they apply to real-life dynamic scenarios are necessary should to clarify this point. In a recent study, we have demonstrated that semantically consistent sounds can speed up search latencies for an object in dynamic and naturalistic visual scenes (Kvasova et al., 2019). This finding proves that audio-visual congruency facilitation effects for task-relevant objects demonstrated with simple and artificial AV stimuli (Experiments 1B and 1C; Iordanescu et al., 2008, 2010) could be generalized to the real-world contexts. However, it is perhaps fair to say that in real-life conditions, most of the sensory information available (including audio-visual congruent objects if present) occur outside the focus of attention and are potentially task irrelevant. Therefore, experiments with realistic scenes should address the effects of cross-modal semantic congruence in task-irrelevant or no-task conditions. These validations in more ecologically valid materials will help understand the relevance of semantic congruence in real life.

## Conclusions

In the current study, we examined the constrains under which audio-visual semantic congruence triggers spatial orienting. We found that audio-visual semantic congruence speeded visual search times when the cross-modal objects are task relevant, a phenomenon that had been already described in other studies. Here, we show that even when these audio-visually congruent objects are task irrelevant they can summon attention, but only when presented under low perceptual load conditions. When these audio-visual events are irrelevant to the task and perceptual load is high, then the attention-grabbing effects of audio-visually congruent events vanish. This pattern of results does not support a strict automaticity hypothesis of semantic integration across modalities. Instead, we believe that some top-down processing is necessary for audio-visual semantic congruence to trigger spatial orienting. Further, in order to understand the relevance of semantic congruence in real life more experiments with realistic scenes should address the cross-modal semantic effects in task-irrelevant or no-task conditions.

## Supporting information

Visual stimuli

## Acknowledgements

This research was supported by the Ministerio de Economia y Competitividad (PSI2016-75558-P AEI/FEDER), AGAUR Generalitat de Catalunya (2017 SGR 1545). Daria Kvasova was supported by an FI scholarship, from the AGAUR Generalitat de Catalunya. This manuscript has been released as a pre-print at bioRxiv (Kvasova and Soto-Faraco, 2019).

1 That is to say. At the moment the task-irrelevant sound-image combination was presented, there was no other competing task or stimuli.

