## Supplementary figures and images for "Not so automatic: Task relevance and perceptual load modulate cross-modal semantic congruence effects on spatial orienting"

### Supplementary.png

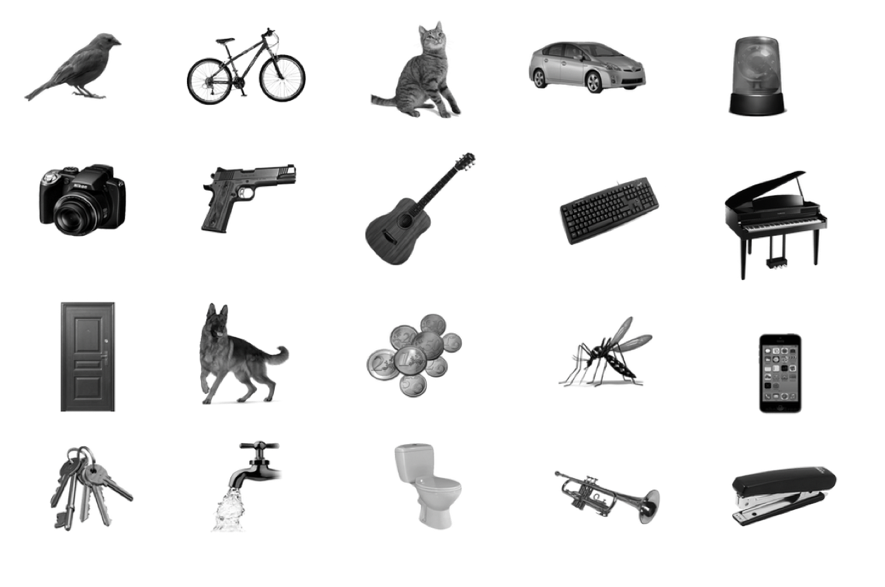
